# Distinct molecular mechanisms shape thermal tolerance and its plasticity in a splash pool copepod

**DOI:** 10.64898/2026.07.29.741598

**Authors:** Rujuta V. Vaidya, Isabelle P. Neylan, Deborah Shombaowo, Brant C. Faircloth, Maheshi Dassanayake, Morgan W. Kelly

**Affiliations:** Department of Biological Sciences, Louisiana State University, Baton Rouge, LA 70803; Museum of Natural Science, Louisiana State University, Baton Rouge, LA 70803

**Keywords:** thermal tolerance, plasticity of heat tolerance, gene expression, trade-offs, heat shock response

## Abstract

Understanding how organisms respond to changes in temperature is becoming increasingly important in our rapidly changing world. While species differ substantially in their thermal tolerance, the exact molecular mechanisms underpinning this trait variation remain largely unknown. Species with broad geographical distribution provide a unique opportunity to examine variable responses to temperature, as populations experience a diverse range of thermal regimes that differ in the intensity, frequency, and duration of thermal stress. We used a splash pool copepod (*Tigriopus californicus*) with a large geographic range across the North American coast (Baja California to Alaska) to examine variation in thermotolerance and the plasticity of thermotolerance in five populations sampled across 12 degrees of latitude. We found that populations with higher heat tolerance (southern) showed lower plasticity (measured as increased survival at higher temperatures after a prior exposure to sub-lethal temperature). Our comparative transcriptomic analyses revealed that higher heat tolerance was associated with maintaining elevated expression of genes coding for peptidases even in absence of heat shock and higher plasticity of heat tolerance was associated with an increase in gene expression plasticity of genes coding for chitin and extracellular matrix (ECM). Our results highlight ontology-specific patterns associated with changes in the trait means and plasticity of heat tolerance across populations. These findings improve our mechanistic knowledge of thermotolerance as a trait and our ability to predict which populations are most vulnerable to extinction based on the trade-off between fixed thermal limits and plastic responses.

## Introduction

A rapidly warming world is exposing organisms to new and unpredictable temperature regimes. Temperature affects life at all scales, from structural integrity of biological structures to the rates of reactions in the cell, to the dynamics and persistence of populations, to ecological interactions within communities (Angilletta Jr. 2009, Hochachka & Somero, 2002, Jawad et al. 2024). Understanding how organisms respond to temperature is therefore essential for understanding current distributions and predicting which species and populations will persist in the future as warming continues (Parmesan 2006; Kearney and Porter 2009; Somero 2010). Most measurements of thermal limits have focused on variation across species; however, species with broad geographic distributions also experience a wide range of temperatures, and distinct populations of the same species often show differences in their heat tolerance (Kuo and Sanford 2009; Eliason et al. 2011; Kelly et al. 2012; Dongmo et al. 2021). This variation in heat tolerance will play an important role in species responses to climate change (Jawad et al. 2026) and also represents an opportunity to investigate the mechanisms associated with variation in heat tolerance.

Intraspecific variation in heat tolerance may arise from genetic differences among individuals, but also from phenotypic plasticity, where a single genotype can give rise to multiple phenotypes (West-Eberhard, 2003; Ghalambor et al. 2015). Phenotypic plasticity can buffer populations from immediate selection pressures and can also serve as the mechanism that drives adaptation (Price et al. 2003, Bell et al. 2009, Chevin et al. 2013, Neylan et al. 2026). In the case of heat tolerance, many species exhibit a form of phenotypic plasticity called ‘heat hardening’, where brief exposure to a stressful but sublethal heat shock confers a short-lived increase in heat tolerance immediately after the initial shock (Precht 1973, Bowler et al. 2005, Bilyk et al. 2012, Earhart et al. 2022). Populations of the same species can differ in their baseline thermal tolerance and potentially also in the plasticity of this baseline tolerance (i.e., their capacity for heat hardening; Sørensen et al. 2003; Tomanek 2010; Zhao et al. 2017;Van Heerwaarden and Kellermann 2020; Barley et al. 2021; Hoffmann et al. 2023).

At the molecular level, one way populations can achieve higher heat tolerance is by maintaining elevated expression levels of genes involved in the heat shock response, even in the absence of heat stress. This pattern, called ‘transcriptional frontloading’ has been documented in many ectotherms, and is proposed as mechanism to explain why populations of the same species can achieve different heat tolerances (Barshis et al. 2013; Fifer et al. 2021; Rivera et al. 2021; Vidal-Dupiol et al. 2022; Brown et al. 2025). Alternatively, higher heat tolerance can also be achieved through higher plasticity of gene expression, such that tolerant populations are better at rapidly and accurately changing their gene expression after experiencing heat stress (Kelly 2019). This gene expression plasticity has been proposed to be a more metabolically efficient alternative to achieve higher tolerance when maintaining higher gene expression levels is energetically demanding (Rivera et al. 2021) (Fig. 1A). These same strategies, i.e. changes to baseline expression levels or changes to gene expression plasticity, could also drive population specific differences in heat hardening ability. After experiencing the initial shock, populations can acquire higher heat tolerance either by continuing to maintain elevated expression levels of genes involved in heat shock response, or by increasing their gene expression plasticity, which can confer improved tolerance during a subsequent shock (Pratx et al. 2024, Georgoulis et al 2021, Oberkofler et al. 2021, Hilker et al. 2016) (Fig. 1B).

**Figure 1:**
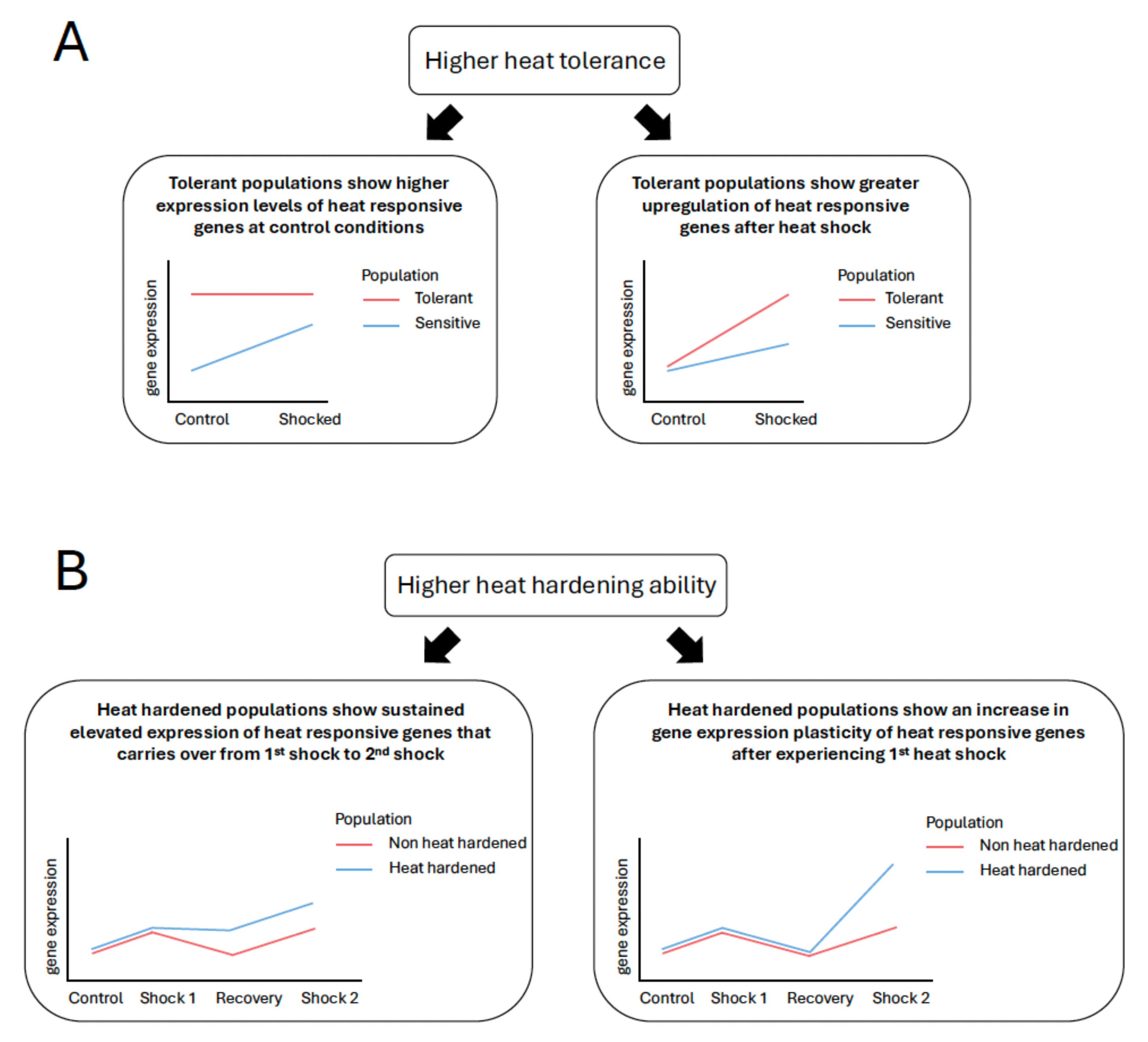
A conceptual schematic overview outlining the hypothetical patterns of gene expression that may explain. A) higher heat tolerance and B) higher heat hardening ability seen within populations of *Tigriopus californicus*.

Here, we investigated the mechanisms associated with variation in heat tolerance and its plasticity among populations of the splashpool copepod, *Tigriopus californicus*. *T. californicus* occupies shallow intertidal pools in a habitat that stretches from Baja California to Alaska along the North American coast, and *T. californicus* populations experiences a wide range of temperature variation over multiple spatial and temporal scales (Burton and Lee 1994; Kelly et al. 2012). As a result, *T. californicus* has evolved strong local adaptation to temperature, with different populations showing significant variation in the temperature they can tolerate as well as their heat hardening capacity (Kelly et al. 2021; Pereira et al. 2017; Barreto et al. 2018; Foley et al. 2019; Bogan et al. 2024). Combined with their short generation time (3-4 weeks), a suite of genomic and transcriptomic resources, and ease of maintenance in the lab, *Tigriopus* has emerged as a promising study system to explore intra-specific variation in responses to temperature and the mechanisms that govern it (Schoville et al. 2012; Kelly et al. 2013, 2017; Tangwancharoen and Burton 2014; Graham and Barreto 2019; Harada et al. 2019; Neylan et al. 2026; Vaidya et al. 2025).

We first measured heat tolerance and plasticity of heat tolerance in five *T. californicus* populations and then compared gene expression profiles to determine. mechanisms associated with higher heat tolerance and higher plasticity of heat tolerance in these populations. If higher heat tolerance in *Tigriopus* populations was achieved through transcriptomic frontloading, we expected tolerant populations to show higher baseline expression of heat-responsive genes at control conditions relative to more sensitive populations. If higher tolerance was driven by higher plasticity of gene expression, we expected tolerant populations would show greater up regulation of heat responsive genes after heat shock compared to sensitive populations. Similarly, if heat hardening is caused by higher baseline expression or greater plasticity of heat responsive genes post heat shock, we expected that populations with greater heat hardening capacity would show sustained elevated gene expression levels of heat responsive genes that carried over from the first shock to the second shock or an increase in gene expression plasticity of heat responsive genes after the first shock. Lastly, we also wanted to test if the mechanisms for achieving higher heat tolerance and higher plasticity are mutually exclusive, so we compared population level differences in gene expression in response to two shocks. By testing these predictions, we hoped to gain insights into the striking variation in thermal tolerance seen in this species and improve our understanding of the cellular mechanisms that shape this plastic trait.

## Methods

### Animal collection and maintenance

We collected *T. californicus* copepods from five sites in California and Oregon, USA. Specifically, between December 2023 to December 2024, we collected our southern populations from Sunset Cliffs (SU, 32° 43’ 56.83“N, –117° 15’ 24.04” W) and Bird Rock (BD: 32°49′ N, – 117°16′ W) in San Diego California and we collected northern populations from Salt Point, California (SA, 38° 20′ N, –123° 33′ W), and Bodega Marine Reserve, California (BR, 38°04′ N, –123°19′ W) in California, and from Strawberry Hill, Oregon (SH, 44°15′ N, –124°06′ W) (Fig. 2A). We established lab cultures from these wild–caught individuals following protocols described in Kelly et al. (2012). Briefly, copepods were maintained at 19°C and at 35 ppt salinity under a 12 hour light / 12 hour dark cycle and fed a diet of ground spirulina fish food *ad libitum*. Each population was maintained in the lab for at least three generations before beginning the experiments.

**Fig 2.**
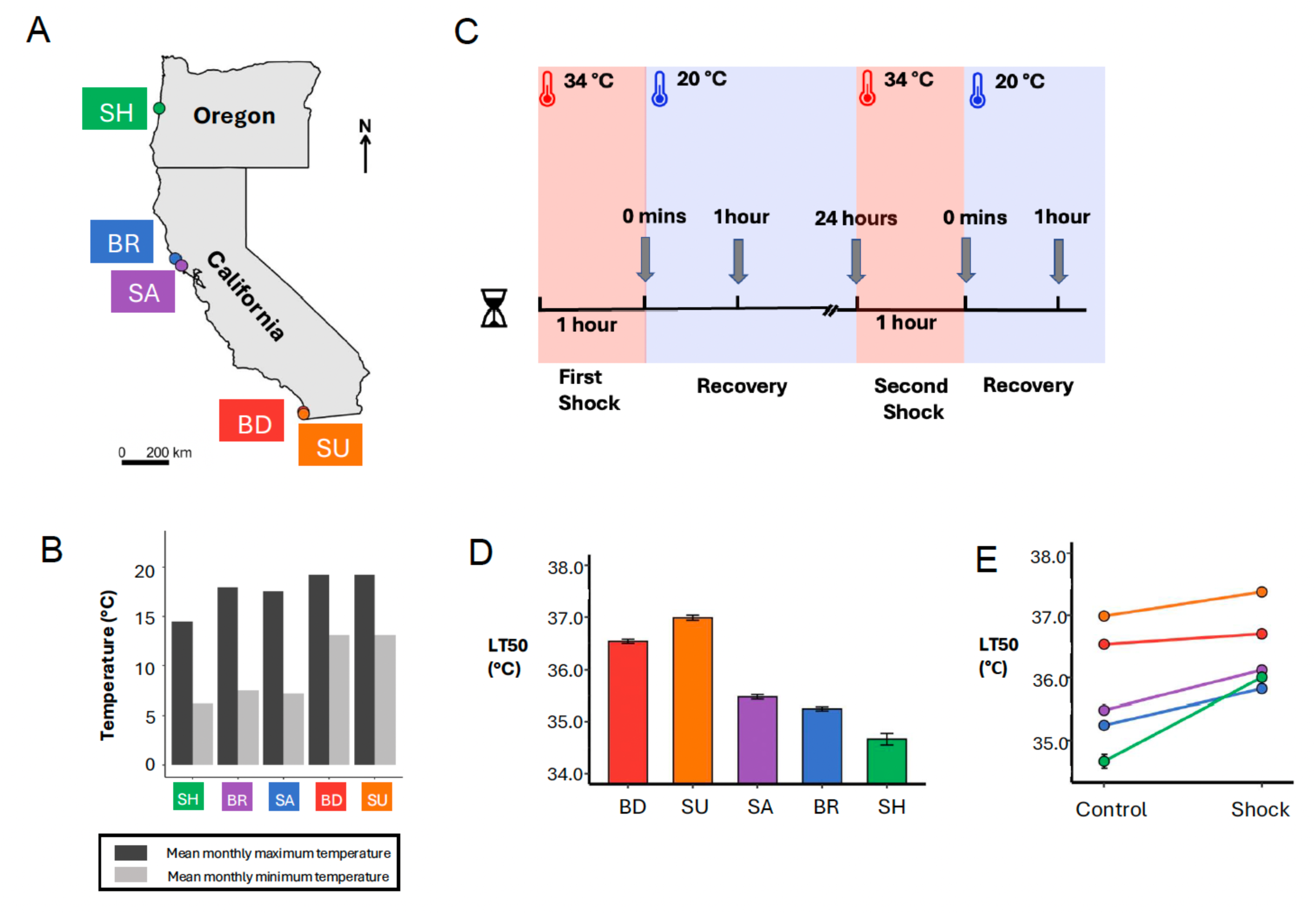
A: Sampling sites for the five *Tigriopus californicus* populations used in this study. Sunset Cliffs, California (SU, 32° 43’ 56.83“N, –117° 15’ 24.04” W), Bird Rock, California (BD: 32°49′ N, –117°16′ W), Salt Point, California (SA, 38° 20′ N, –123° 33′ W), Bodega Marine Reserve, California (BR, 38°04′ N, –123°19′ W), Strawberry Hill, Oregon (SH, 44°15′ N, –124°06′ W). B: Monthly minimum and maximum temperatures (°C) for our five collection sites, according to the data extracted from Worldclim. Populations are denoted on x-axis and values on y-axis represent temperature in °C. C. A schematic design of our transcriptomic experiment showing heat shocks in red and recovery periods in blue. Grey arrows indicate sampling points at which animals were flash frozen for RNA extraction. D. The results of the baseline heat tolerance assays for these populations for all five populations. Values along the y-axis represent the temperature (°C) at which 50% of the animals died (LT50) and population names are indicated across the x-axis. Error bars represent standard error. Southern populations are represented by warmer colors (BD: red, SU: orange) and northern populations are represented by cooler colors (SA: purple, BR: blue, SH: green). E. A reaction norm style plot showing measurements of plasticity of heat tolerance (heat hardening ability) for these five populations, measured as the change in their LT50 value after experiencing a prior sub-lethal heat shock. The x-axis indicates treatment groups: control (no prior sub lethal shock before measuring LT50) and shock (animals received a sub-lethal shock prior to LT50 measurements). Values on y-axis represent the temperature (°C) at which 50% of the animals died (LT50), and error bars indicate standard error. The color scheme for the line plots is same as 2.A.

### Measurements of thermal tolerance and plasticity of thermal tolerance

To measure thermal tolerance and thermal tolerance plasticity, we subdivided each population into three lines and maintained these lines separately throughout the experiment to give us three biological replicates per population. We measured thermal tolerance by estimating LT50 temperature values, or the temperature at which 50% of the animals die (Kelly et al 2012). To estimate the LT50 values, we exposed copepods to a set of temperatures ranging from temperatures that showed 100% survival to 100% mortality. We began our assays at 34.5°C, which was 0.5°C higher than their sub-lethal shock and marked our 100% survival datapoint.

Then we continued to expose individual sets of copepods to a range of temperatures increasing in steps of 0.2°C until we reached a temperature that showed 100% mortality. Each temperature was tested with at least three replicates. Each replicate consisted of three mate-guarded copepod pairs that we haphazardly chose from one of our three lines for each population. We placed these six copepods into a 0.6mL PCR tube filled with 200uL of artificial sea water at 35 ppt salinity and then placed the tubes into a thermocycler (Techne 3PRIMEX/02, Bibby Scientific LTD) to precisely control temperature. In the thermocycler, copepods were subjected to 1 hour acclimation period at 30°C, before ramping up to the target temperature, where they were held for another hour. Copepods were allowed to recover for 24 hours, after which we counted the number of surviving copepods in each tube. We defined heat hardening as an increase in thermal tolerance (measured as an LT50 value) after prior exposure to a sub-lethal heat shock. To estimate heat hardening, we first shocked mate-guarded pairs from each population at a sub-lethal temperature of 34 °C for 1 hour in a water bath (VWR WBE20 General Purpose Water Bath) (Vaidya et al 2025, Neylan et al 2026, Kelly et al, 2012). The animals were allowed to recover for 24 hours, after which we tested their heat tolerance by following the same protocol above. We used the drc package (Ritz, 2015) to calculate LT50 temperatures. For each population, we fit a logistic regression model (LL.2) that allowed us to estimate an LT50 value as well as the standard error associated with this estimate for our baseline heat tolerance and plasticity of heat tolerance measurements. All analyses were performed in R version 4.4.0 (R Core Team, 2023).

### Experimental design for transcriptomic analyses

For our transcriptomic experiment, we used a different subset of copepods originating from the same lines and culture stocks that we used for our phenotypic measurements for our transcriptomic experiment. We performed two sub-lethal heat shocks to closely mimic our phenotypic measurements of baseline heat tolerance and plasticity. Our experimental design (Fig. 2C) consisted of three time points following the first heat shock that we used to identify gene expression patterns associated with differences in baseline heat tolerance. We included two timepoints after a subsequent second shock to identify the gene expression patterns associated with the plasticity of heat tolerance. Because heat hardening occurs within a few hours of the initial heat shock, we also included these multiple timepoints after first and second shock to help us compare gene expression changes over time across the two shocks. Each time point consisted of three replicates, and we used one non-heat shocked control treatment for each of our replicate lines within populations. For each population, we haphazardly selected 50–60 male and female adult copepods (copepods past their final molting stage) from the established lab cultures and transferred these into a 50 mL Falcon© tube containing 50 mL of sea water at 35 ppt salinity. Each time point (and the non-shocked control) had three Falcon© tubes containing 50-60 copepods, which served as three biological replicates for that specific experimental treatment (3 Falcon© tubes for non-shocked controls, 3 Falcon© tubes for each of the five heat-shocked timepoints for a total of 18 falcon tubes in total for each population). For the first heat shock, we placed these tubes in a water bath (VWR WBE20 General Purpose Water Bath) for 1 hour at 34°C. After the shock, copepods were returned to the room temperature (about 22°C) where they were maintained until they were flash frozen in liquid nitrogen at their respective time points following the heat shock (0 minutes, 1 hour, and 24 hours after shock) (Fig. 2C). A subset of these tubes from the first shock were allowed to recover for 24 hours and were shocked again (our second sub-lethal shock) following the same protocol as the first shock. We flash froze copepods at 0 minutes and 1 hour after this second shock. The control tubes were kept at room temperature for the duration of heat shock and were all flash frozen by the end of the 24 hours after our first heat shock. The flash frozen samples were stored in –80°C until RNA extractions.

### RNA extraction and sequencing

We extracted total RNA from flash frozen copepods using a combination of TRIzol reagent (Invitrogen; catalog no. 15596026) and the Qiagen RNAeasy Plus kit (Qiagen, catalog no. 74134), following a previously described protocol (Neylan and Vaidya 2025). Briefly, copepods were homogenized using a TissueRuptor II (Qiagen, catalog no. 9002755) before we followed steps 1–7 of the TRIzol protocol, followed by steps 4–11 of RNAeasy Plus kit protocol. Total RNA extracted from 90 samples (((5 timepoints + 1 control) x 3 replicates) x 5 populations) were sent to Novogene Corporation Inc. at Sacramento, California, where RNA quality was confirmed using a 2100 Agilent Bioanalyzer on a Eukaryote Total RNA Nano chip and non– directional libraries were produced using poly–A tail selection. The resulting 90 libraries were sequenced on NovaSeq X Plus, with 150bp paired end reads. We removed adapter sequences using Trimmomatic (Bolger et al. 2014), and we used the program FASTQC (Andrews, 2010) to ensure that all of the reads considered for downstream analysis had quality scores of at least 35 (Table SX). The reads were then mapped to the *T. californicus* reference genome (Barreto et al. 2018; NCBI GCF_007210705.1) using the STAR aligner (version 2.6.0a; Dobin et al., 2013). Reads were mapped to gene features with the options (––quantMode GeneCounts –– outFilterScoreMinOverLread 0.50 ––outFilterMatchNminOverLread 0.50) to adjust for poly–A tail contamination (Sirovy et al. 2021), which generates a count matrix using ReadsPerGene.out.tab output. Transcripts per million (TPM) values used for visualizing our RNASeq dataset were generated using the RSEM package (version 1.3.3; Li and Dewey 2011). All downstream analyses were performed in R version 4.0.3 software (R Core Team 2021).

### Differential gene expression analysis

We removed lowly expressed genes, which we defined as genes that did not have at least 10 counts in 25% of our samples (Love et al. 2014, DESeq2 vignette) from our dataset. We then used our filtered datasets as input to DESeq2 (v 1.24.0) (Love et al. 2014) to perform pairwise comparisons of each time point relative to controls and to obtain a list of differentially expressed genes (DEGs) for each time point. The false discovery rates (FDRs) were calculated using the Benjamini–Hochberg method (Benjamini and Hochberg 1995). Genes with an adjusted p–value cutoff of < 0.05 and a log2foldchange of > 1 (for upregulated genes) or < –1 (for downregulated genes) were considered to be differentially expressed. Gene ontology (GO) enrichment for each comparison was performed using customized TopGO (citation) scripts and the Fisher’s Exact Test (p < 0.05).

### Analysis of elevated baseline gene expression in tolerant populations

To test if the higher baseline heat tolerance was achieved through maintaining higher baseline expression levels (in absence of heat shock) of genes involved in heat shock response, we combined DEGs upregulated in response to the first and the second heat shock for all populations at all timepoints. Then removed duplicates from this list to build a consensus list of 1837 genes (hereafter referred to as “heat responsive genes”). We then extracted TPM values for these heat responsive genes from the three control replicates for each population. We calculated z-scores for the expression of each gene across all replicates and all populations before averaging z scores for replicate values within a given population. A positive z-score for a given gene in a given population indicated a gene was expressed at a higher baseline level (at control conditions, in absence of heat shock) in that population compared to the others. If higher heat tolerance is produced through higher baseline expression of heat responsive genes (Fig 1A) then we expected a greater number of heat responsive genes with greater baseline expression under control conditions in the more heat tolerant populations. For each population, we also performed gene ontology (GO) enrichment analyses for genes that showed higher baseline using custom TopGO (Alexa and Rahnenführer, 2026) scripts and Fisher’s Exact Test (p < 0.05).

### Analysis of plasticity of gene expression across first and second shock

To visualize gene expression plasticity, we constructed gene expression reaction norms to examine changes in gene expression (TPM values) in our five populations across two heat shocks. Briefly, for each population we computed average TPM values (averaged across three replicates) for heat responsive genes across our experimental timepoints and visualized the output as line plots using ggplot (Wickham 2011). Differential gene expression can be visualized as the slopes of lines (represented by TPM averages) between control and shocked timepoints. A positive slope indicated an increase in gene expression and a negative in slope indicated a decrease in gene expression (Rivera et al 2021). We also used these line plots to examine the changes in magnitude of gene expression across two shocks in our five populations. Using lme4 package in R (Bates et al, 2015) to run linear mixed-effects model, we tested whether the magnitude of upregulation after a heat shock (visualized as the slopes of the lines in our line plots) differed between southern and northern populations across first and second shocks for our gene categories of interest.

## Results

### A) Thermal tolerance assays reveal that populations with higher heat tolerance show lower plasticity of heat tolerance

We observed higher baseline heat tolerance in the southern populations than the northern populations. Among all five populations, SU had the highest (36.99 ±0.04), and SH had the lowest (34.66 ±0.11) heat tolerance (Fig. 2D). We also found that overall, southern populations had lower plasticity of heat tolerance than the northern populations. Our northernmost population SH was most plastic (an increase of 1.03°C in LT50), followed by BR (an increase of 0.64°C in LT50), SA (an increase of 0.59°C in LT50), SU (an increase of 0.39°C in LT50), and BD (an increase of 0.19°C in LT50) (Fig. 2E).

### Higher heat tolerance of southern populations is associated with elevated baseline expression of peptidases and higher gene expression plasticity of Heat Shock Proteins

Our comparison of baseline gene expression of 1837 heat responsive genes (Table S2) revealed that under control conditions, the more heat tolerant southern populations (SU and BD) had a greater number of genes showing higher baseline expression levels than the less heat tolerant northern populations (Fig. 3B and 3C). SU had the most (1248) heat responsive genes showing higher baseline expression across all five populations, while BD had 631 genes with higher baseline expression. However, our northernmost population SH, was an exception to our hypotheses. SH showed the second highest number of genes (906 genes) with higher baseline expression. The remaining two northern populations (BR and SA) ranked lowest, with 625 and 491 genes respectively. We also visualized the average TPM values of these 1837 heat responsive genes across our experimental timepoints as gene expression reaction norms to look for population-specific patterns (Fig.4A). We observed higher gene expression plasticity (visualized as slopes of the gene expression reaction norms in Fig. 4A) in our heat tolerant, southern populations than northern populations. One of our northern populations (BR) was an exception to trend and showed equally higher gene expression plasticity as our southern population (BD).

**Figure 3.**
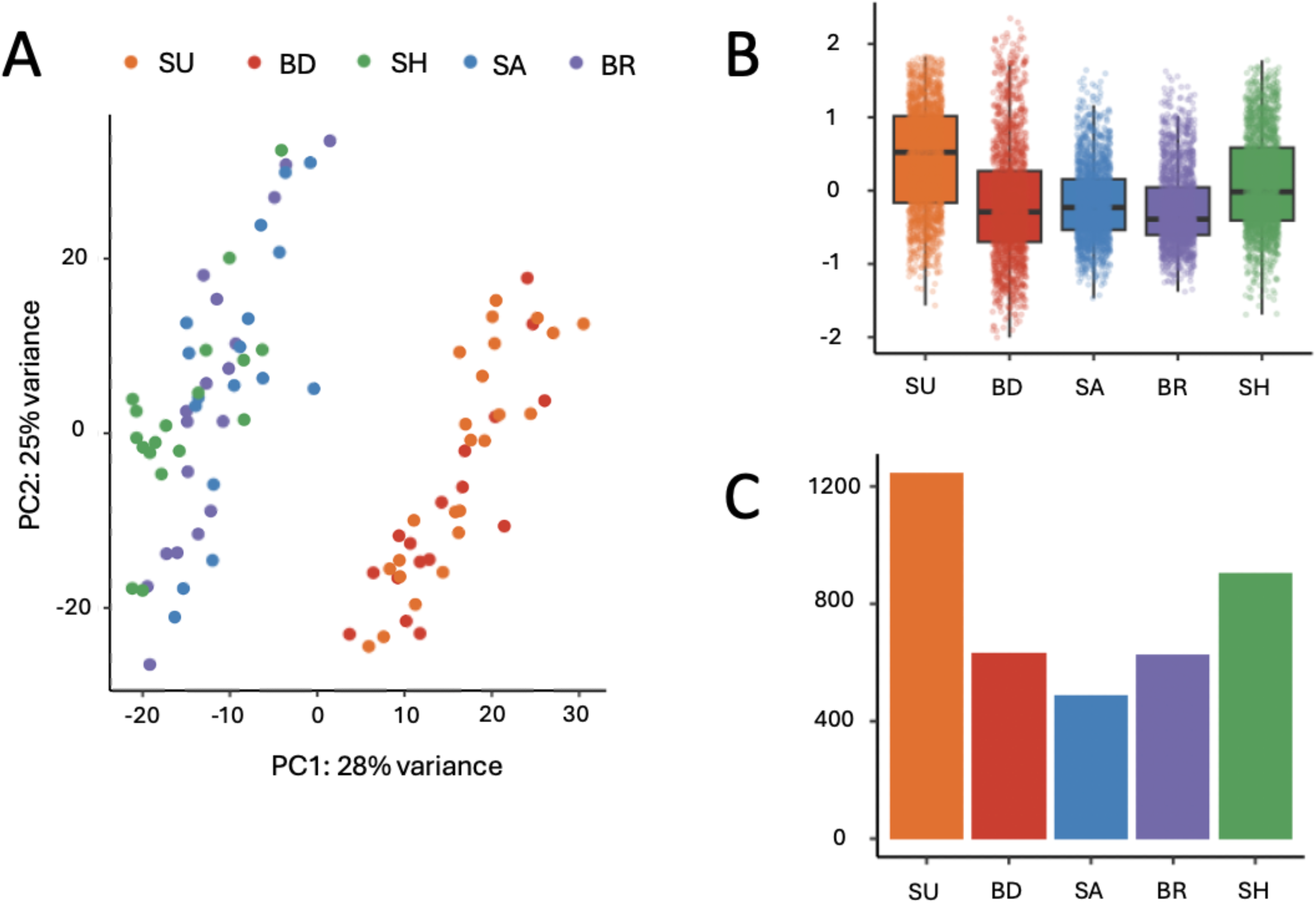
A: Principal component analysis (PCA) plot for all five populations generated using plotPCA function of DESeq2 (top 500 most variable genes). B: Boxplots showing mean z scores of TPM values for heat responsive genes under control conditions across all five populations. Populations are denoted on X axis, while the Y axis shows z score values. C: Bar plots showing total number of heat responsive genes that showed higher baseline expression levels in each population. Y axis indicates number of genes and populations are denoted on X axis.

**Figure 4:**
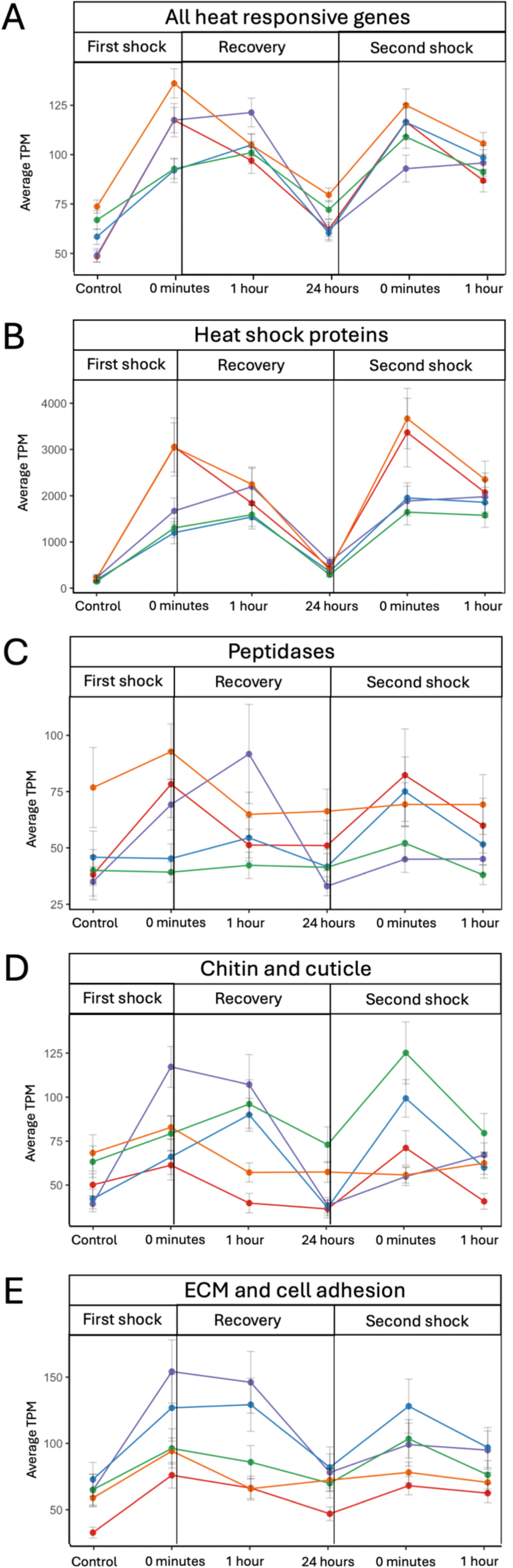
Line plots showing gene expression levels (mean transcript per million (TPM) values; Y axis) for all heat responsive and their specific gene categories their sub-categories across first (A) and second (B) shock. Y axis for the first shock indicates control and 0 minutes timepoints collected after first heat shock, while Y axis for the second shock indicates 24 hours after heat shock and 0 minutes timepoints collected after second heat shock. Line colors indicate populations: SU (orange), BD (red), SH (green), BR (purple), SA (blue). Error bars indicate standard error values for mean TPMs.

Prompted by these unexpected gene expression patterns, we also evaluated whether populations differed in identities of the genes that showed elevated baseline expression levels at control conditions and higher gene expression plasticity after the first shock. From our 1837 heat responsive genes, we visualized gene encoding: heat shock proteins, peptidases, chitin and cuticle component, and extracellular matrix and cell adhesion (Fig. 4B-E). We observed that genes coding for heat shock proteins showed higher upregulation after the first shock in southern populations than northern populations (detected through average TPM values and visualized as differences in slopes of the gene expression reaction norms in Fig. 4B). We found that genes coding for peptidases showed elevated baseline expression, but lower gene expression plasticity in our most heat tolerant population (SU) and interestingly also showed lower gene expression plasticity in two of our northern, less tolerant populations (SA and SH) (Fig. 4C). On the other hand, genes coding for chitin and cuticle elements were notably elevated at baseline in our most heat sensitive population (SH) along with the two tolerant southern populations but showed highest plasticity of gene expression in one of our less heat tolerant populations (BR) (Fig. 4D). Finally, we detected higher baseline expression levels as well as gene expression plasticity for genes coding for elements of cell matrix and adhesion in northern populations relative to southern populations (Fig. 4E).

### Higher plasticity of heat tolerance is associated with sustained elevated expression levels and higher gene expression plasticity of genes involved in structural integrity

To test whether the first shock induced a pattern of elevated gene expression that carried over beyond the shock, we compared average TPM values of heat responsive genes at control and 24 hours post heat shock. We observed that the average TPM values of our 1837 heat responsive genes were higher in all five populations relative to their controls, indicating a pattern of sustained elevated gene expression that continued even after the shock (Fig. 4A). To assess the impact of a subsequent shock on gene expression, we treated our 24 hour timepoint after the first shock as our control for the second shock.

Beyond these global patterns across of heat responsive genes, we also observed gene category specific expression patterns in response to second shock across our five populations. We observed sustained elevated expression levels of genes coding for HSPs, peptidases, and cell matrix and adhesion at 24 hours after first shock in the southern populations (Fig. 4 B-E). On the other hand, we observed a decrease in gene expression plasticity of genes coding for peptidases (*p*=0.0001), cell matrix and adhesion (*p* < 0.0001), and chitin and cuticle ( *p* < 0.0003) (visualized as slopes of the gene expression reaction norms in Fig. 4 B-E) in southern populations when compared with northern populations which showed an increase in gene expression plasticity for those genes. Additionally, we also observed sustained elevated expression levels at 24 hours after first shock for genes encoding chitin and cuticle components (SH) and cell matrix and adhesion (SA, BR, SH) in our northern populations.

## Discussion

We were interested in understanding the molecular mechanisms governing variation in heat tolerance and its plasticity in five distinct populations of *T. californicus*. We predicted that higher heat tolerance could be achieved either through maintaining higher baseline expression levels of heat responsive genes or through higher gene expression plasticity. We also expected that populations with higher plasticity of heat tolerance would show sustained elevated expression levels of heat responsive genes or higher gene expression plasticity that improved tolerance during a subsequent heat shock. We found evidence for both of these predicted mechanisms: elevated gene expression (either at baseline or induced after initial heat shock) and transcriptional plasticity emerged as shared mechanisms underlying both higher heat tolerance and higher plasticity. However, the functional categories of genes driving these patterns were distinct, pointing to separate molecular mechanisms for achieving tolerance versus plasticity in this system. In our phenotypic data, we found evidence of a trade-off between plasticity and baseline tolerance with more heat tolerant populations showing lower plasticity. This trade-off pattern has been widely observed in metazoans including *Tigriopus* (Barley et al 2021, Neylan et al 2025, Kelly et al 2017, Brennan et al 2022, Sasaki et al 2021, Bogan et al 2024).

Our transcriptomic data revealed that the greater heat tolerance in the southern populations was associated with higher baseline expression (peptidases) and greater plasticity of gene expression (HSPs). HSPs and peptidases are key components of the heat shock response (HSR), which is a highly conserved cellular defense mechanism that protects cells from heat stress induced damage (Morimoto 1998; Feder and Hofmann 1999; Sørensen et al. 2003; Richter et al. 2010; De Nadal et al. 2011). At a cellular level, heat stress compromises protein structural stability, causing denaturation and loss of function. The resulting accumulation of cytotoxic protein aggregates can be fatal to a cell, necessitating prompt clearance (Hochachka and Somero 2002; Stefani and Dobson 2003). HSPs mitigate this damage by binding to denatured proteins to prevent their further aggregation (Hartl et al. 2011; Chen et al. 2018), whereas peptidases clear existing aggregates by degrading irreversibly damaged proteins (Lipscomb 1980; Rawlings and Bateman 2019). Notably, while HSPs typically serve as an inducible defense mechanism, peptidases can be preemptively expressed and maintained in the cell (Sørensen et al, 2003, Richter et al, 2010, Kidrič et al, 2014). This dual strategy of leveraging inducible/plastic HSPs and maintaining elevated baseline peptidase expressions suggests that southern populations achieved higher thermal tolerance through modifying distinct components of the heat shock response.

The lower baseline expression levels of peptidases, and lower plasticity for HSPs we observed in the northern populations suggests at an absence of similar investment in counteracting protein damage caused by heat stress, which is reflected in their lower heat tolerance. This pattern of populations achieving higher thermal tolerance through maintaining higher gene expression levels of heat responsive genes pre-emptively, in absence of thermal stress, has been well-documented in other marine taxa (Barshis et al. 2013; Fifer et al. 2021;

Rivera et al. 2021; Vidal-Dupiol et al. 2022; Brown et al. 2025).

The association between HSPs and peptidases focused gene expression changes and higher heat tolerance was further underscored in our most sensitive northern populations, SH. Despite having the second-highest number of genes with elevated baseline expression, SH exhibited the lowest baseline heat tolerance. Interestingly, the specific genes that were elevated at baseline in SH were predominantly associated with chitin metabolism, extracellular matrix structure, and cellular adhesion; while our analyses indicated that expression of these genes was responsive to heat shock, these genes are not classically considered components of the heat shock response. Although structural components like chitin and collagen are vital for exoskeleton and cellular integrity (Schoville et al. 2012; Mouw et al. 2014; Graham and Barreto 2019; Harada et al. 2019; Winkler et al. 2020; Chakravarti et al. 2022; Matsubayashi 2022; Andriot 2024), their higher baseline expression did not confer higher heat tolerance in the SH population. Taken together with the lower gene expression plasticity of HSPs that we observed in the SH population, our findings demonstrated that achieving higher heat tolerance in *Tigriopus* requires modifications to components of the heat shock response rather than structural stability of the cell.

The lower heat hardening capacity observed in the southern populations was marked by expression levels of HSPs that remained similar across both shocks while genes coding for peptidases showed a decrease in gene expression plasticity after receiving the second shock. This diminished response across the southern populations we sampled could stem from having sufficient amounts of peptidases already transcribed in the cell or they could result from due to a potential physiological ceiling effect (Barua and Heckathorn 2004; Somero 2010; Tomanek 2010). We also observed a similar loss of plasticity of gene expression in genes encoding for cell matrix and adhesion, chitin and cuticle, and lipid transport and metabolism in our southern populations in response to the second shock. Upregulation of genes involved in cuticle and extracellular matrix has been associated with remodeling the heat induced structural damage inside the cell in *T. californicus* as well as in other marine taxa (Tomanek and Somero 1999; Watabe 2002; Dong et al. 2008; Mouw et al., 2014; Barreto 2019; Harada et al. 2019; Andriot 2024). The reduced plasticity of these structural genes suggests that the heat stress response in southern populations is largely focused on HSPs and peptidases, and this cellular investment in components of heat shock response consequently results in a loss of gene expression plasticity for several other genes affected by changes in temperature. While this cellular strategy has enabled them to tolerate higher temperatures, the loss of plasticity in a large subset of genes responding to heat shock following a second shock could potentially limit their ability to sense and adapt their cellular response to thermal stress. It is also possible that the southern populations have reached the tipping point of their metabolic budget, as maintaining higher baseline expression levels could be metabolically expensive (Rivera et al. 2021; Alagar Boopathy et al. 2022). Coupled with the higher baseline levels of these energetically demanding proteins, southern populations may not have enough energetic budget left to invest in upregulation of genes involved in cell structure and integrity, and rebuild the structural damage caused by heat stress.

Interestingly, we see the exact opposite pattern in the northern populations we sampled, where the second shock increased the gene expression plasticity of genes encoding for chitin and cuticle components, extracellular matrix, cell adhesion, and lipid transport and metabolism. These expression patterns suggest that in contrast to their southern counterparts, northern populations have achieved higher plasticity of heat tolerance via mechanisms that allow for sensing and fine-tuning the cellular response to thermal stress across a broad range of biological processes including protein damage and maintaining cellular integrity. Taken together, our gene expression comparisons suggest that distinct cellular mechanisms are associated with higher plasticity and higher tolerance in this system. Future studies that investigate other fitness and physiological costs, as well as occurrences of mRNA and protein modifications in response to thermal stress will certainly reveal a more complete picture of thermal adaptation in *T. californicus*.

Our findings highlight how populations of the same species that experience different temperature regimes can vary in their cellular responses to thermal stress. With climate change altering both the frequency and intensity of severe heat events, understanding how populations may differ in their abilities to tolerate thermal stress is essential for predicting their continued survival. Our results show that while northern populations have the physiological and cellular ability to adapt and increase their baseline thermal tolerance, the southern populations appear to be approaching their physiological limit. These stark population level contrasts within a species highlight the need to account for population level differences in thermal tolerance for predicting current and future persistence estimates for species in the face of climate change.

## Author Contributions

RVV, MWK conceived the ideas and designed methodology; IPN, DS, and RVV collected the data; IPN and RVV analyzed the data; RVV and MWK led the writing of the manuscript; BCF, IPN, and MD contributed critically to revisions; All authors reviewed the paper and gave final approval for publication.

## Acknowledgements

We thank Zaiah Herbert, Elizabeth Heneghan, Christian Mack, and Emma Crable for their assistance in copepod care. We thank Wissam Jawad, Sofía Savicki-Kelly, Jonah Savicki-Kelly, Akhil Bhardwaj, and Pratik Barge for their help with copepod collections in the field. Portions of this research were conducted with high performance computing resources provided by Louisiana State University (http://www.hpc.lsu.edu). Funding was provided by the National Science Foundation through grant IOS 2154283 (to MWK, BCF, MD) and a Postdoctoral Research Fellowship in Biology 2305966 (to IPN).

## Data availability statement

Sequencing data associated with this study will be deposited on NCBI and will be available after manuscript is accepted for publication.

## Supplementary tables

Supplementary table S1: Number of significantly differential expressed genes (DEGs) detected for each pairwise comparison performed using DESeq2.

Supplementary table S2: Comparison of differences in expression levels (TPM) under control conditions among five populations.

Supplementary table S3: TPM values (averaged across three replicates) for each experimental timepoint and gene category, listed for all five populations sampled in our study.

Supplementary Table S4: List of significantly differentially expressed genes (calculated via DESeq2 pairwise comparisons) in control (non-heat shocked) vs heat shocked copepods from BR population.

Supplementary Table S5: List of significantly differentially expressed genes (calculated via DESeq2 pairwise comparisons) in control (non-heat shocked) vs heat shocked copepods from SA population.

Supplementary Table S6: List of significantly differentially expressed genes (calculated via DESeq2 pairwise comparisons) in control (non-heat shocked) vs heat shocked copepods from BD population.

Supplementary Table S7: List of significantly differentially expressed genes (calculated via DESeq2 pairwise comparisons) in control (non-heat shocked) vs heat shocked copepods from SU population.

Supplementary Table S8: List of significantly differentially expressed genes (calculated via DESeq2 pairwise comparisons) in control (non-heat shocked) vs heat shocked copepods from SH population.

Supplementary table S9: Gene Ontology terms overrepresented in significantly differentially expressed genes in control vs heat shocked copepods from BR population.

Supplementary table S10: Gene Ontology terms overrepresented in significantly differentially expressed genes in control vs heat shocked copepods from SA population.

Supplementary table S11: Gene Ontology terms overrepresented in significantly differentially expressed genes in control vs heat shocked copepods from SH population.

Supplementary table S12: Gene Ontology terms overrepresented in significantly differentially expressed genes in control vs heat shocked copepods from BD population.

Supplementary table S13: Gene Ontology terms overrepresented in significantly differentially expressed genes in control vs heat shocked copepods from SU population.

